# Amino acid substitutions in norovirus VP1 dictate cell tropism via an attachment process dependent on membrane mobility

**DOI:** 10.1101/2023.02.17.528071

**Authors:** Jake T. Mills, Susanna C. Minogue, Joseph S. Snowden, Wynter K.C. Arden, David J. Rowlands, Nicola J. Stonehouse, Christiane E. Wobus, Morgan R. Herod

**Affiliations:** Astbury Centre for Structural Molecular Biology, School of Molecular & Cellular Biology, Faculty of Biological Sciences, University of Leeds, Leeds, UK; Department of Microbiology and Immunology, University of Michigan Medical School, Ann Arbor, MI 48130, USA

**Keywords:** MNV, murine norovirus, CD300lf, receptor, membrane fluidity, virus evolution

## Abstract

Viruses interact with receptors on the cell surface to initiate and co-ordinate infection. The distribution of receptors on host cells can be a key determinant of viral tropism and host infection. Unravelling the complex nature of virus-receptor interactions is, therefore, of fundamental importance to understanding viral pathogenesis. Noroviruses are non-enveloped, icosahedral, positive-sense RNA viruses of global importance to human health, with no approved vaccine or antiviral agent available. Here we use murine norovirus as a model for the study of molecular mechanisms of virus-receptor interactions. We show that variation at a single amino acid residue in the major viral capsid protein had a key impact on the interaction between virus and receptor. This variation did not affect virion production or virus growth kinetics, but a specific amino acid was rapidly selected through evolution experiments, and significantly improved cellular attachment when infecting immune cells in suspension. However, reducing plasma membrane mobility counteracted this phenotype, providing insight into for the role of membrane fluidity and receptor recruitment in norovirus cellular attachment. When the infectivity of a panel of recombinant viruses with single amino acid variations was compared *in vivo*, there were significant differences in the distribution of viruses in a murine model, demonstrating a role in cellular tropism *in vivo*. Overall, these results highlight the importance of lipid rafts and virus-induced receptor recruitment in viral infection, as well as how capsid evolution can greatly influence cellular tropism, within-host spread and pathogenicity.

**Importance:** All viruses initiate infection by utilising receptors to attach to target host cells. These virus-receptor interactions can therefore dictate viral replication and pathogenesis. Understanding the nature of virus-receptor interactions could also be important to developing novel therapies. Noroviruses are non-enveloped icosahedral viruses of medical importance. They are a common cause of acute gastroenteritis with no approved vaccine or therapy and are a tractable model for studying fundamental virus biology. In this study, we utilise the murine norovirus model system to show that variation in a single amino acid of the major capsid protein can alone can affect viral infectivity through improved attachment to suspension cells. Reducing plasma membrane mobility reduced infectivity, providing an insight into the importance of membrane mobility for receptor recruitment. Furthermore, variation at this site was able to change viral distribution in a murine model, illustrating how in-host capsid evolution can influence viral infectivity and immune evasion.

## Introduction

Cellular tropism is a key determinant for viral infection of a host and is dictated by several factors, including viral attachment to cellular receptors. Unravelling the complex nature of virus-receptor interactions is therefore of fundamental importance to understanding viral pathogenesis. Human noroviruses (HNV) cause gastroenteritis and are responsible for >200,000 deaths and a cost of ∼£40 billion worldwide each year (1). With no efficacious vaccine or approved therapy to treat HNV infections, a greater understanding of the virus life cycle and capsid structure is likely to be important for developing new approaches to disease control.

Noroviruses are members of the *Caliciviridae* family of positive-sense single-stranded RNA viruses (1), that have three or four open reading frames (ORF) 1-4 (2). ORF1 is translated to produce the viral polyprotein that is cleaved to generate the non-structural (NS) proteins required for genome replication (2). ORF2 and 3 encode the two viral structural proteins, VP1 and VP2, respectively (2). ORF4 is only expressed in murine norovirus (MNV) and encodes virulence factor 1 (VF1) (3). The two viral structural proteins assemble to enclose the genome in a *T = 3* capsid. This protein shell is ∼40 nm in diameter and is composed of 180 copies (90 dimers) of the major structural protein VP1, and a low copy number of the minor structural protein VP2 (4). In feline calicivirus, 12 copies of VP2 forms a portal-like assembly likely involved in genome release, but this is yet to be demonstrated for other caliciviruses (4). VP1 monomers comprise an N-terminal region, a shell (S) domain, and a protruding (P) domain. The P domain is additionally split into the proximal and distal sub-domains, P1 and P2, respectively (5, 6). *In vitro* replication of HNV has been demonstrated in human intestinal enteroids (7), human B cells (8) and salivary gland cells (9), but these models are technically challenging, highly variable (10), and suffer from the lack of an effective reverse genetics system. Consequently, MNV is frequently used as a model system for the study of norovirus structure and pathology.

MNV is widely prevalent in laboratory mice (11). MNV-1 was the first strain of MNV to be identified (12), and it establishes acute, self-resolving infections in wild-type mice, but can be fatal in immune compromised (STAT1^-/-^) mice (13). Different strains of MNV have different cellular tropisms, which in turn determine the site(s) of infection in the host. Strains such as MNV-3 are located primarily in the colon and caecum (14), while MNV-1 is detected across the gastrointestinal tract and in immune cells (15), including macrophages and dendritic cells, thought to aid virus distribution to extra-intestinal sites (16–18). Furthermore, MNV-3 can still be detected in the faeces 56 days post-infection and can establish lifelong persistent infections (14). This draws parallels with HNV infection, whereby virus shedding can be detected up to 28 days post-infection (19), and persistent infection in immunocompromised individuals can last years (20). Cellular tropism is also important in determining MNV persistence, with serotypes such as MNV-CR6 able to infect rare tuft cells located in the intestinal epithelium and evade the immune system (21, 22).

The cellular tropism of MNV is thought to be determined by expression of CD300lf, the primary proteinaceous receptor (with the virus also able to utilise CD300ld to enter the cell) (24, 25). Both CD300lf and CD300ld are members of the CD300 receptor family of type I transmembrane proteins with a 2 disulphide bond extracellular domain (26). They are present on numerous immune cell types such as dendritic cells, where they are thought to play opposing roles to maintain homeostasis (24, 27). Since the identification of CD300lf as the physiological receptor for MNV (28), studies have begun to dissect the nature of this interaction. The P2 sub-domain of the VP1 capsid directly interacts with the receptor, with two CD300lf ectodomains binding one P2 sub-domain (25). The interaction mimics the way phospholipids bind to the receptor, is conserved across multiple MNV serotypes, and is enhanced by divalent cations (Ca^2+^ and Mg^2+^) and bile acid (25, 29, 30). Structural studies have suggested that up to 21 amino acids of VP1 form a network of interactions with 19 residues of CD300lf (5, 24, 25). Despite this extensive network of interactions, the binding affinity is reported to be low (KD:∼219 μM), therefore, receptor avidity may be important for endocytosis (25). Studies have attempted to elucidate how genetic variation in VP1 can influence cellular tropism and pathogenesis (23, 31–33), however more research is needed.

Using the MNV model system, we demonstrate that variation in a single amino acid in the major capsid protein can alter virus-receptor interactions in cell culture, as well as within-host spread in the mouse model. Specifically, our experiments suggest that a single substitution at this site can enhance cell specific growth in culture by allowing more robust recruitment of multiple receptors under conditions of high membrane fluidity. Consistent with this idea, reducing membrane mobility significantly reduced viral infection. Finally, this amino acid variation affects tissue tropism in mice, which has implications for within-host spread and organ-specific infection. Together, these results reveal information on how viruses utilise membrane fluidity to overcome low-affinity receptor interactions, and how the plasticity of the viral capsid can affect cellular and organ tropism.

## Results

### Identification of key residues in VP1 for MNV infectivity

Previous studies identified 21 amino acids of MNV VP1 that form a network of interactions with the receptor CD300lf (24, 25, 34). Through alignment of all available MNV sequences, most of these residues are highly or completely conserved across MNV isolates, however, one residue, VP1 301, showed considerable variability (Figure 1A). Furthermore, we noted that there was an association between the residue encoded in this position and viral strains, i.e. MNV-1 and MNV-4 predominantly encode threonine (T) whilst all other strains predominantly encode isoleucine (I). We therefore set out to investigate how variations in the identity of this VP1 residue could influence viral replication and pathogenesis.

**Figure 1:**
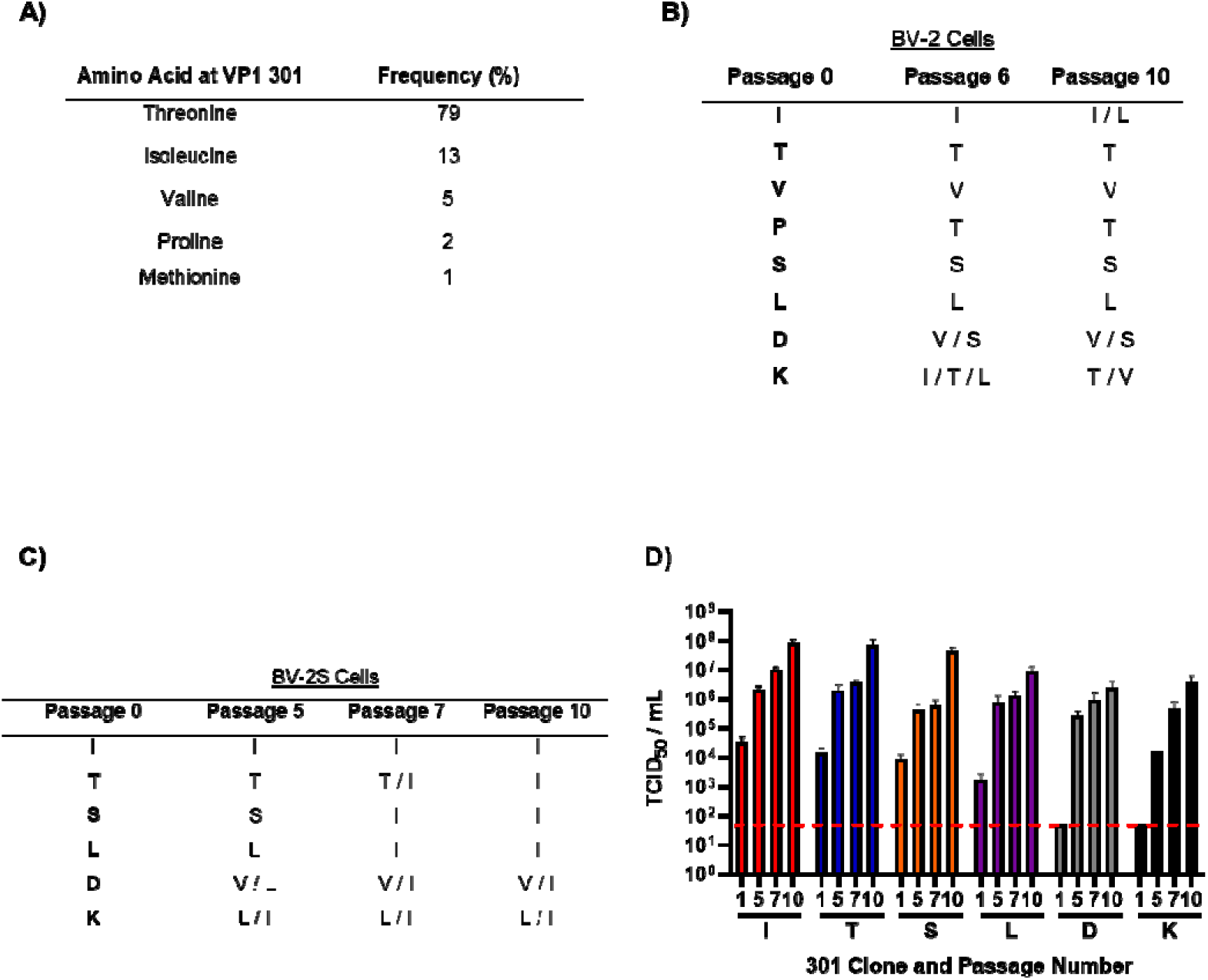
Repeat passaging of MNV-1 in suspension leads to selection of hydrophobic residues at VP1 301. **(A)** Overall amino acid variation at MNV VP1 301 was plotted from deposited sequences on GenBank. Recombinant MNV-1.CW1 with single amino acid substitutions in the infectious clones were passaged 10 times in **(B)** adherent BV-2 cells or **(C)** BV2 cells in suspension (BV-2S), before the ORF2 was sequenced at indicated passages. Data shows amino acid residues encoded at the position 301 of VP1 (n = 3). **(D)** Viruses passaged through BV-2S cells were titrated at selected passages as indicated. Red dotted line demonstrates limit of detection for TCID_50_ assay. Data shows mean TCID_50_/mL (n = 3 ± SEM).

We began by investigating whether variants at this amino acid position were genetically stable through cell culture evolution experiments. To ensure a homogenous genetic background, we modified an infectious clone of MNV-1.CW1 (that encodes T at amino acid 301 of VP1), to encode either, I, valine (V), or proline (P). All of these amino acids have been documented at this position in MNV sequences deposited to GenBank. In order to ascertain the importance of the amino acid at this position, we also generated infectious clones with serine (S), leucine (L), aspartic acid (D) or lysine (K). These infectious clones were used to produce *in vitro* transcribed RNA and virus was recovered by transfection of BHK-21 cells (termed passage 0). The recovered viruses were serially passaged 10 times in BV-2 cells grown adherently or in suspension (for brevity termed BV-2S). RNA was extracted from virus samples taken at indicated passages, reverse transcribed, and the consensus ORF2 sequence determined (Figure 1B and C).

When passaged in adherent BV-2 cells (Figure 1B), the VP1 sequence for the MNV-1.CW1 infectious clones carrying T301, V301, S301 and L301 did not change throughout the experiment. With the I301 infectious clone, two out of the three replicates maintained I301 at passage 10, whilst an I301L substitution occurred in the third replicate by passage 10. MNV-1.CW1 infectious clones carrying P301, D301 and K301 all underwent substitution at this position by passage 6 to encode a range of amino acids, which narrowed by passage 10 to P301T, D301V/S and K301T/V. For all of the sequences there were no other amino acid changes throughout ORF2.

In BV-2S cells, only the MNV-1.CW1 I301 infectious clone was stable and did not acquire any VP1 amino acid substitutions throughout the experiment (Figure 1C). In contrast, substitutions were found in MNV-1.CW1 T301, S301 and L301 infectious clones to encode isoleucine at the consensus level (T301I, S301I and L301I) between passage 5 and 7 (Figure 1C). Again, in infectious clones of MNV-1.CW1 encoding D301 or K301, substitution of D301V/I and K301L/T in the consensus sequence was detected by passage 5 (Figure 1C). Importantly, there were no other changes to the wild-type sequence of MNV-1.CW1 VP1 in any of the infectious clones. To determine the effects of these substitutions on viral yield, supernatants from the BV-2S passage experiment were titrated by TCID_50_ assay on BV-2 cells (Figure 1D).

The titre of the MNV-1.CW1 I301 clone increased over the duration of the experiment from ∼1×10^4^ TCID_50_/mL at passage 1 to ∼1×10^8^ TCID_50_/mL by passage 10, which was the peak titre for any virus. Infectious clones carrying MNV-1.CW1 T301, S301, and L301 (that all changed to 301I) followed a similar pattern, having initial titres between 1×10^3^ - 1×10^4^ TCID_50_/mL, before increasing to ∼1×10^7^ TCID_50_/mL by passage 10. The infectivity of MNV-1.CW1 D301 and K301 were below the limit of detection (LOD) until passage 5, when the titre increased to ∼1×10^4^ and _∼_1×10^5^ TCID_50_/mL, respectively, before the titre reached a peak of ∼1×10^7^ TCID_50_/mL at passage 10. This increase in titre coincided with the change to hydrophobic residues, with a preference for 301I. Together, these data suggest that viruses with isoleucine at VP1 position 301 have a particular advantage when grown in suspension cell culture.

### The VP1 301 amino acid is a major determinant for infectious virus production in suspension cultures

To confirm that VP1 I301 conferred increased viral infectivity in suspension cell culture, the virus yield following transfection of BHK-21 cells with RNA was determined in BV-2 cells. RNA transcribed *in vitro* from the infectious clones was transfected into BHK-21 cells which are permissive for viral replication but do not express the viral receptor, therefore the amount of infectious virus detected is directly proportional to the replication of the transfected RNA alone. Virus was collected and titrated by TCID_50_ assays on suspension grown BV-2S cells (Figure 2A), adherently grown BV-2 cells (Figure 2B) or BV-2 cells grown adherently but infected in suspension (Figure 2C). For suspension TCID_50_ assays, viral dilutions were prepared and added to the plates first, before cells were seeded.

**Figure 2:**
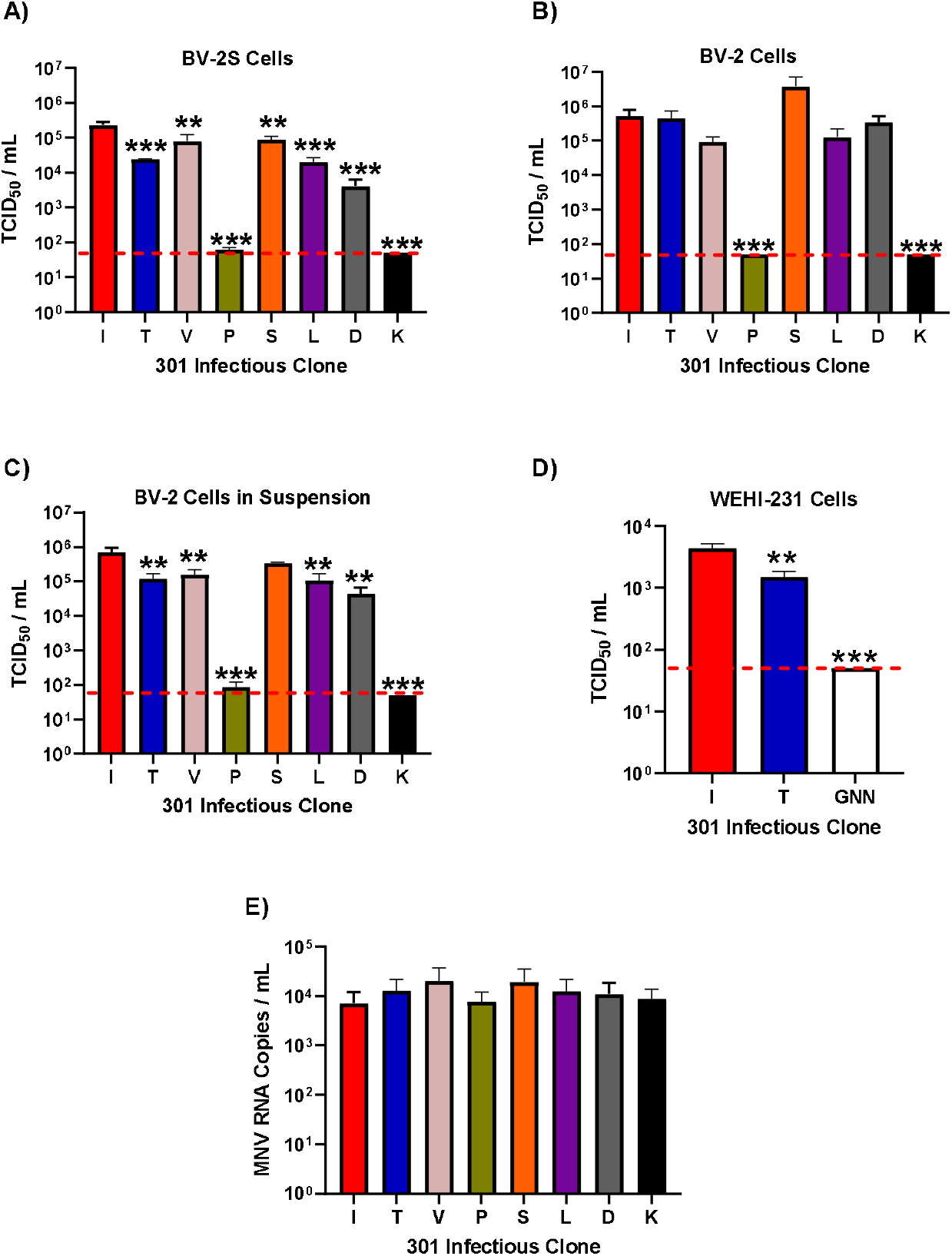
The VP1 301 residue is a major determinant of virus particle infectivity *in vitr*o. MNV-1.CW1 infectious clone RNAs with the indicated amino acids at VP1 301 were transfected into BHK-21 cells and virus-containing supernatants collected after 48 hours. Virus titre was determined by TCID_50_ assays on **(A)** suspension BV-2S cells, **(B)** adherent BV-2 cells, **(C)** adherent BV-2 cells infected in suspension or **(D)** WEHI-231 B lymphocyte suspension cells. The WEHI-231 experiment also contained an RdRp replication-defective MNV GNN negative control. Data shows mean TCID_50_/mL, with significance compared to I301 using one-way ANOVA with corrections for multiple comparisons (n = 3 ± SEM, *p<0.05; **p<0.01; ***p<0.001). The red dotted line demonstrates limit of detection for TCID_50_ assays. **(E)** In separate transfections, the recovered supernatant was treated with 25 U/mL benzonase for 30 minutes at 37□C, before RNA was extracted. RNA concentration was then measured by one-step RT-qPCR. Data show mean RNA genome copies/mL (n = 3 ± SEM).

On BV-2S cells (Figure 2A), the titre of the MNV-1.CW1 I301 variant was significantly higher than all other infectious clones. This was ∼5-fold higher than MNV-1.CW1 V301 and S301 and ∼10-fold greater than MNV-1.CW1 T301, L301 and D301. Both MNV-1.CW1 P301 and K301 variants had titres below the LOD, suggesting these substitutions are detrimental to MNV infectivity.

In contrast, when the infectious clones were titrated on adherently grown BV2 cells (Figure 2B), there were no significant differences in the titre of MNV-1.CW1 S301, I301, T301, V301, L301 and D301 viruses, with titres all between 1×10^5^ TCID_50_/mL and 1×10^6^ TCID_50_/mL. Again, MNV-1.CW1 P301 and K301 were highly detrimental for infectivity.

To understand whether this observation was specific for cells grown or infected in suspension, the TCID_50_ assays were repeated with adherently grown BV-2 cells, however, the infection was performed while the cells were in suspension before being allowed to adhere to the culture vessels (Figure 2C). In this setup, the titre of the MNV-1.CW1 I301 variant was again significantly higher than all other infectious clones, except MNV-1.CW1 S301, with both titres ∼10-fold greater than for MNV-1.CW1 T301, V301, L301 and D301 variants. Once again, MNV-1.CW1 P301 and K301 viral recovery was at or below the LOD.

To rule out differences in transfection efficiency, we conducted similar experiments whereby select infectious clone RNA was co-transfected into BHK-21 cells alongside an IRES-GFP DNA plasmid. The measurement of GFP fluorescence alongside titration of the recovered virus allowed us to correct the viral titre for variation in transfection efficiency. Following collection, the virus was titred by TCID_50_ assay in the same three cell infection conditions and normalised to GFP fluorescence at 24 hours post-transfection.

Once again, MNV-1.CW1 I301 had a significantly greater viral titre compared to all other infectious clones when the TCID_50_ assay was conducted in BV-2S cells (Supplemental Figure 1A). There was no significant difference in viral titres between infectious clones in adherent BV-2 cells (Supplemental Figure 1B). Although there was no significant difference in adherent BV-2 cells infected in suspension (Supplemental Figure 1C), there was a similar pattern to BV-2S cells across the infectious clones.

Taken together, our data suggest that the MNV-1.CW1 I301 variant has a selective advantage at infecting cells when in suspension, but no selective advantage is observed in adherent cells. To determine whether the differences between MNV-1.CW1 I301 and MNV-1.CW1 T301 viruses applied to another cell type, cell culture infectivity assays were performed in the suspension-grown mouse B lymphocyte cell line WEHI-231. Cells were infected as before and an MTS assay was used to determine cell viability and thus calculate virus infectivity (TCID_50_ assays could not be performed as WEHI-231 cells do not adhere to tissue culture plates; Figure 2D). The viral titre of MNV-1.CW1 I301 was ∼5-fold greater compared to MNV-1.CW1 T301. In comparison, an infectious clone carrying a replication-defective mutation in the viral polymerase (GNN) (35) had viral recovery below the LOD.

One possible explanation for our observations is that some of the VP1 301 variations affect virion assembly, not receptor engagement. To investigate this, the total production of viral particles was measured by one step RT-qPCR. Virus was produced from infectious clone RNA by transfection into BHK-21 cells as before, and non-encapsidated RNA was degraded by nuclease treatment before RNA was extracted and the protected RNA concentration measured by one-step RT-qPCR (Figure 2E). There were no statistically significant differences in the number of virus particles produced by any of the clones.

Taken together, these data indicate that the VP1 301 residue plays an important role in MNV infection, but that this is unrelated to viral replication.

### The amino acid at VP1 301 affects cell attachment

VP1 residue 301 contributes to the virus-CD300lf receptor interface (25) and our data suggested that variation in this amino acid alone is sufficient to confer a replicative advantage to the virus. We hypothesised that the VP1 I301 variant has greater affinity for the receptor, thus increasing cell attachment. To investigate this hypothesis, we conducted virus binding assays with the MNV-1.CW1 I301 or T301 variants on BV-2S cells. We expected that the MNV-1.CW1 I301 variant would bind more effectively to cells in suspension compared to viruses encoding hydrophilic residues.

To prevent endocytosis, binding assays were conducted on BV-2S cells treated with dynasore (Ds), an inhibitor of dynamin that is required for MNV internalisation (36). BV-2S cells were pre-treated with Ds at 37□C, or left untreated as a control, prior to incubation with MNV-1.CW1 I301 or T301 viruses. The amount of virus attached to the cells was measured by western blot for the major viral structural protein, VP1. When analysed by western blot (Figure 3A) and normalised to GAPDH expression, significantly more MNV-1.CW1 I301 binding was detected compared to MNV-1.CW1 T301 in the Ds pre-treated cells (Figure 3B). There was also a trend of increased binding of MNV-1.CW1 I301 compared to MNV-1.CW1 T301 in untreated cells, but this was not statistically significant. BSA was used as a loading control for supernatant, due to the presence of FCS (which contains BSA) in the cell media.

**Figure 3:**
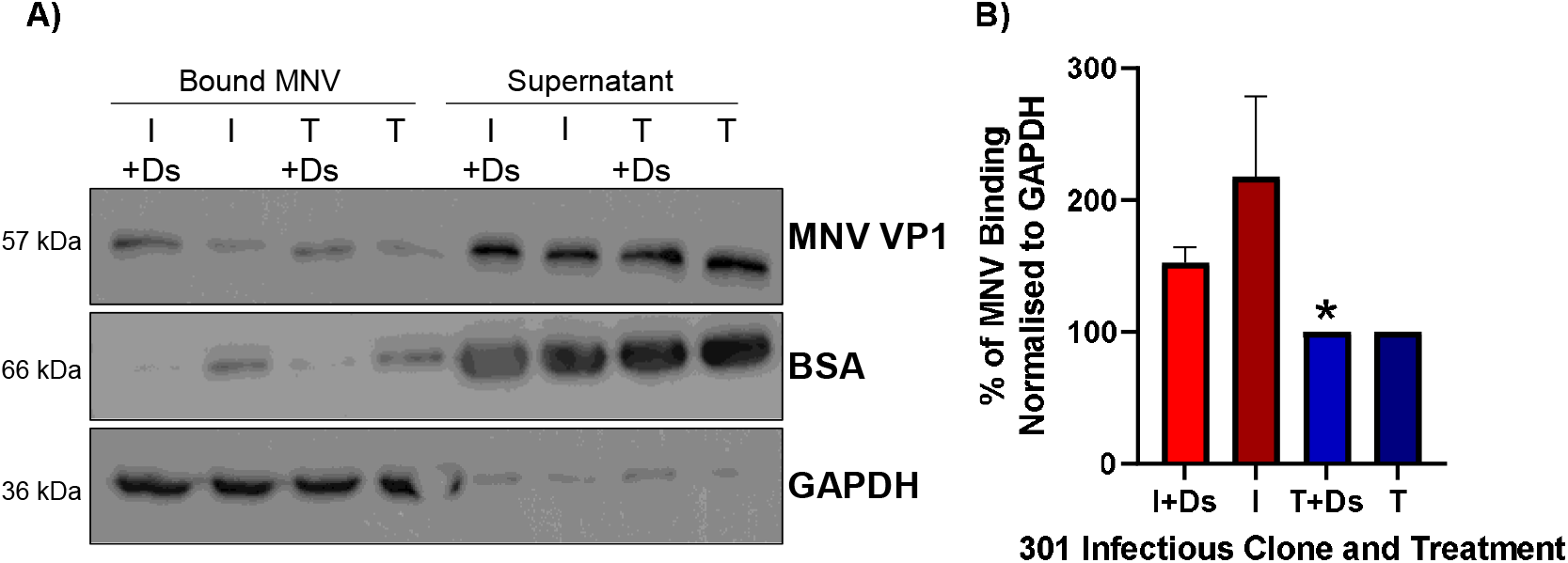
MNV I301 has greater binding capacity than MNV T301 to BV-2S cells. **(A)** BV-2S cells were untreated or pre-incubated with 50 μM dynasore (Ds) for 30 minutes at 37□C, before MNV-1.CW1 I301 or MNV-1.CW1 T301 (MOI 1) was added and incubated for 2 hours at 37□C. Supernatant was removed, cell pellet washed in ice cold PBS before lysis in RIPA buffer. The amount of MNV present in each fraction was quantified by western blot with GAPDH and BSA used as cell-associated and supernatant loading controls, respectively (one representative blot shown). **(B)** The amount of VP1 was quantified by densitometry and normalised to GAPDH. Data shows mean percentage increase/decrease for bound MNV-1.CW1 I301 compared to MNV-1.CW1 T301, with significant differences in virus-cell interaction between MNV-1.CW1 I301 and MNV-1.CW1 T301 infectious clones determined using unpaired T-test (n = 3 ± SEM, *p<0.05).

### MNV binding is dependent on membrane fluidity

Taken together, our observations suggest that viruses encoding I301 have a selective growth advantage and increased cell attachment when in suspension. To confirm that differences between cell types were not due to differences in replication rates, a one-step growth curve with MNV-1.CW1 T301 (10 PFU/cell) was carried out on BV-2 and BV-2S cells. At 12 hours post-infection, there appeared to be an increase in MNV titre in BV-2S cells (albeit not statistically significant), but at the 24 and 48 hour time-points, there was no difference in viral titre (Supplemental Figure 2A). To address potential differences in CD300lf expression between the cell types, we compared the expression levels of CD300lf receptor in BV-2 and BV-2S cells by western blot (Supplemental Figure 2B). The western blot indicated the presence of multiple CD300lf glycosylation states (75 kDa main isoform), as has been previously described (37, 38), however there was no clear difference in relative expression of these different forms between BV-2 and BV-2S cells. BHK-21 cells were used as a negative control, and L-cells and RAW 264.7 cells were used as positive controls for CD300lf expression. Flow cytometry was also used to calculate CD300lf expression in BV-2 and BV-2S cells (Supplemental Figure 2C). There was no difference in CD300lf receptor expression between adherent or suspension cells, with a clear decrease in fluorescence in the controls with no secondary antibody.

Finally, as suspension cells have a greater plasma membrane fluidity compared to adherent cells (39, 40), we hypothesised that fluidity of the cell membrane may affect MNV binding to the cell. We therefore first reduced membrane mobility by performing the binding assay at a reduced temperature. BV-2S cells were pre-treated with Ds at 37□C (to inhibit internalisation), prior to incubation with MNV-1.CW1 I301 or T301 viruses at either 0□C or 37□C. At reduced temperature there was significantly less binding of MNV-1.CW1 to cells, such that little or no VP1 could be detected (Figure 4A & 4B). Next, we treated cells with methyl-β-cyclodextrin (MβCD) and losartan (Los), two compounds reported to chemically restrict membrane mobility (41, 42). To conduct this experiment, BV-2S cells were pre-treated with either Ds, MβCD, Ds & MβCD, or left untreated as a control (Figure 4C & 4D) at 37□C, and either Ds, Los, Ds & Los, or left untreated as a control (Figure 4E & 4F), prior to incubation with MNV-1.CW1 I301 or T301 viruses. MβCD (Figure 4C & 4D) and Los (Figure 4E & 4F) both significantly reduced MNV binding in BV-2S cells by approximately 50% and 30%, respectively. Together, the data indicate the importance of plasma membrane fluidity in MNV cell binding.

**Figure 4:**
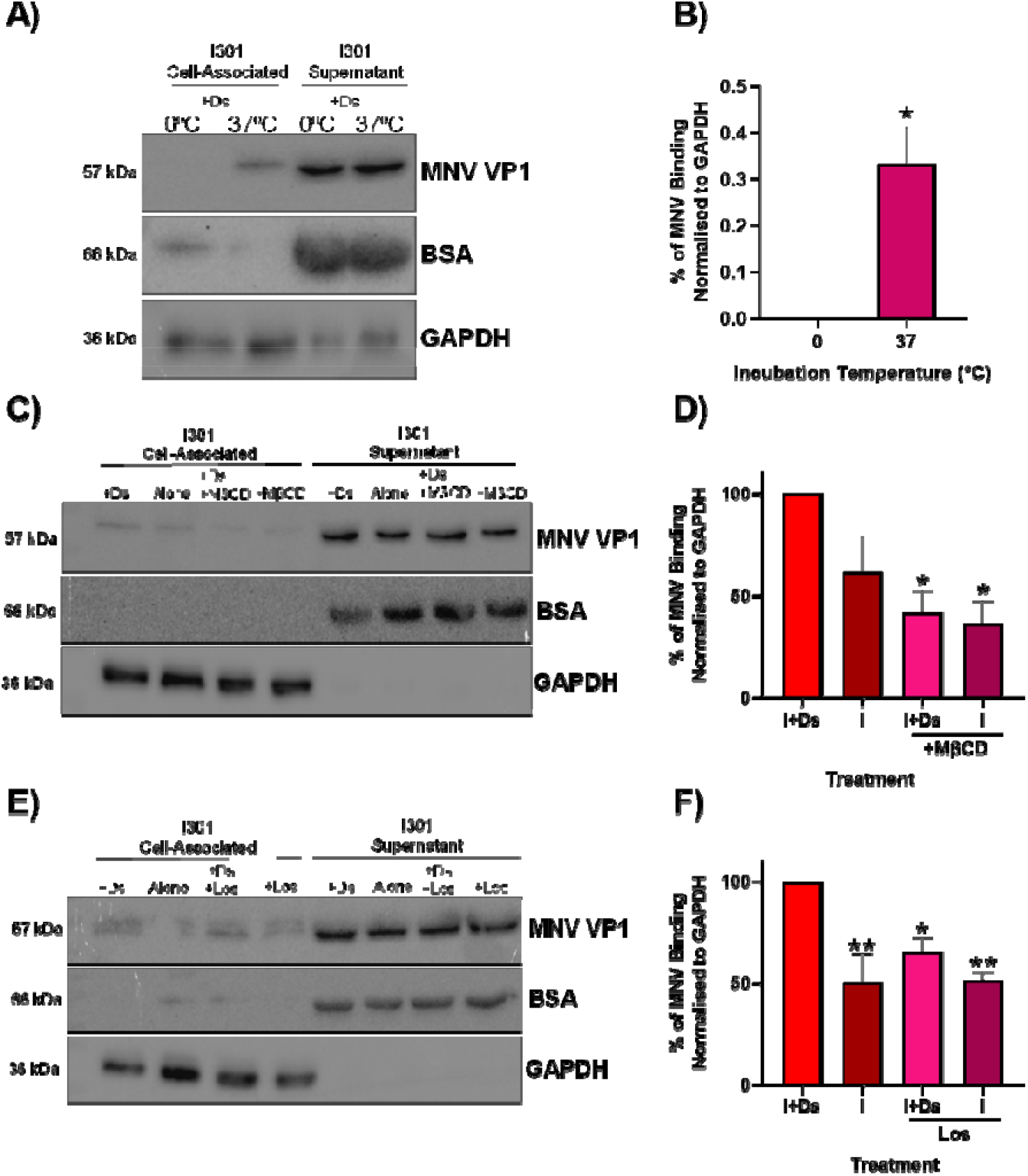
MNV cell binding is temperature dependent and requires membrane fluidity. **(A)** BV-2S cells were pre-incubated with 50 μM dynasore (Ds) for 30 minutes at 37□C, before MNV-1.CW1 I301 (MOI 1) was added and incubated for 2 hours at 0□C or 37□C. Supernatant was removed, cell pellet washed in ice-cold PBS before lysis in RIPA buffer. The amount of MNV present in each fraction was quantified by western blot with GAPDH and BSA used as cell-associated and supernatant loading controls, respectively (one representative blot shown; n = 3). **(B)** VP1 was quantified by densitometry and normalised to GAPDH. Data shows mean densitometry for bound MNV at 0□C and 37□C, with significant differences in virus-cell interaction between temperatures determined using paired T-test (n = 3 ± SEM, *p<0.05). BV-2S cells were untreated or pre-incubated with **(C)** 50 μM Ds, 2 mM methyl-β-cyclodextrin (MβCD) or both Ds and MβCD for up to 60 minutes at 37□C; or **(E)** 50 μM Ds, 40 mM losartan (Los) or both Ds and Los for 30 minutes at 37□C, before MNV-1.CW1 I301 (MOI 1) was added and incubated for 2 hours at 37□C. The binding assay was then performed as in panel A (one representative blot shown; n = 3). The amount of VP1 found was quantified by densitometry and normalised to GAPDH. Data shows mean densitometry for bound MNV, with significant differences in virus-cell interaction between cells **(D)** with or without MβCD pre-treatment; or **(F)** with or without Los pre-treatment determined using one way ANOVA with corrections for multiple comparisons (n = 3 ± SEM, *p<0.05; **p<0.01.)

### The amino acid at VP1 301 influences MNV infectivity and tropism *in vivo*

Our data suggested that MNV-1.CW1 I301 infected suspension cells more effectively than viruses with other amino acids at this position. We therefore hypothesised that this variation at VP1 301 may affect the cellular tropism in a murine model, due to improved infection of non-adherent immune cells at the early stages of infection. To investigate this, seven-week-old C57BL/6 mice were infected by oral gavage with 3×10^5^ PFU/mouse of MNV-1.CW1 I301 or T301. Tissues were harvested from the jejunum (JE), duodenum (DU), ileum (IL), caecum (CE), spleen (SP) and mesentery lymph nodes (ML) at 12 hours post-infection and MNV titre measured by plaque assay (Figure 5).

**Figure 5:**
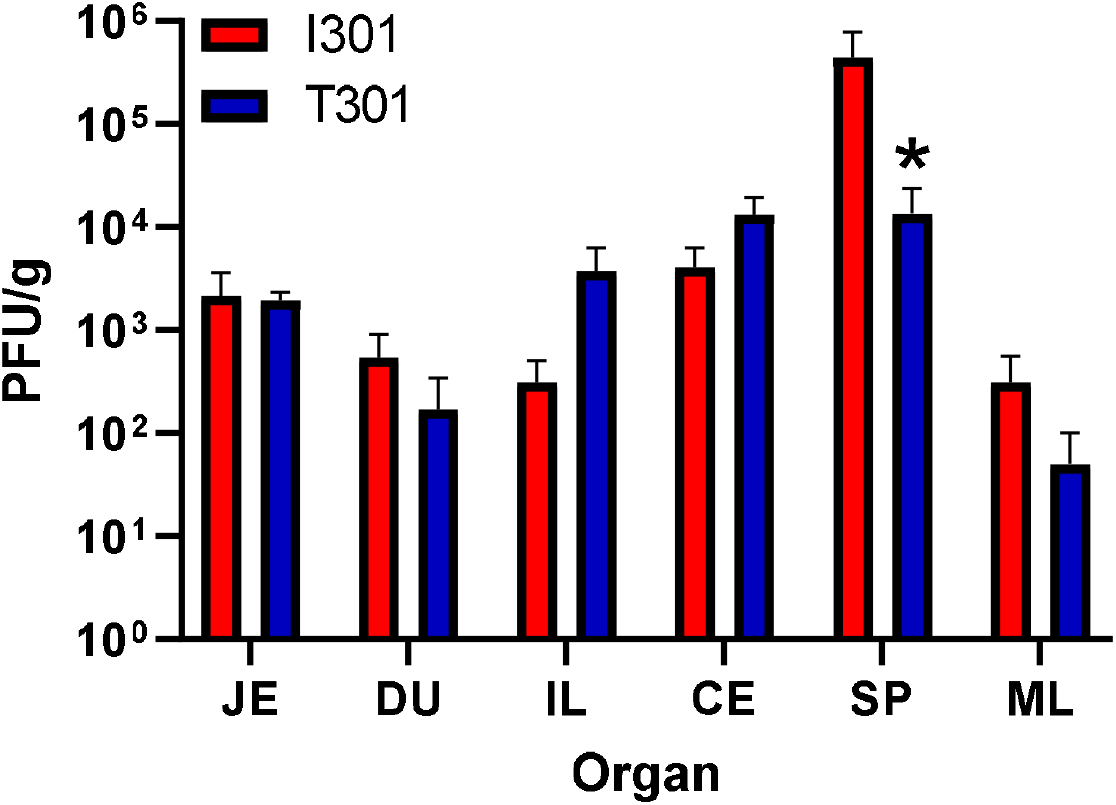
The VP1 301 residue is a determinant of tissue tropism in mice. Seven-week-old C57BL/6 mice were infected by oral gavage with 3×10^5^ PFU/mouse of MNV-1.CW1 I301 or MNV-1.CW1 T301. Mice were sacrificed at 12 hours post-infection, tissues were harvested, and viral titres were determined by plaque assay from the jejunum (JE), duodenum (DU), ileum (IL), caecum (CE), spleen (SP) and mesentery lymph nodes (ML). Plaque numbers (PFU) were normalized to tissue weight (in g). Data show mean PFU/g with significance compared to I301 using two-way ANOVA with corrections for multiple comparisons (n = 5 ± SEM, *p<0.05).

Both MNV-1.CW1 I301 and T301 were detected in all organs after 12 hours of infection. The MNV-1.CW1 I301 viral titre in the SP was significantly higher compared to MNV-1.CW1 T301. There was also a trend of increased MNV-1.CW1 I301 in the ML compared to MNV-1.CW1 T301, and increased MNV-1.CW1 T301 in the IL in comparison to MNV-1.CW1 I301, but these were not statistically significant. To confirm the virus capsid had not undergone mutations during the mouse infection experiments, RNA was extracted from the SP and the sequence of ORF2 determined from the recovered virus. There were no changes identified to the ORF2 consensus sequence extracted from any of the recovered virus compared to the input viruses used for infection. The mouse data agree with our cell culture experiments and suggest that the single amino acid substitution from T to I at position 301 in VP1 can lead to changes in tissue tropism and viral dissemination in the native host.

## Discussion

MNV has a tropism *in vivo* for adherent intestinal epithelial cells, as well as immune cells (13), which are largely non-adherent. Despite this, MNV infection assays in cell culture are usually conducted on adherent macrophage-like cells, ignoring any consequences of infection of cells in suspension. Our work demonstrates that culture and passage of MNV-1 in suspension selects for a single amino acid substitution at VP1 301 that significantly increases infectivity, due to enhanced binding to BV-2S cells. Furthermore, VP1 301 variations are important for the production of infectious virions. The experiments in mice reported here complement the cell culture data, with our findings demonstrating that MNV-1.CW1 I301 has increased cellular tropism in the spleen. Together, this work reveals the significance of the VP1 301 residue in MNV infectivity and pathogenesis.

Norovirus enters the gastrointestinal tract through transcystosis by M cells that are present at the epithelium of Peyer’s patches and at the tips of intestinal villi (43), and once inside the gastrointestinal tract, viruses come into contact with immune cells, such as dendritic cells and macrophages. Our data suggests a mechanism whereby MNV-1.CW1 I301 has greater attachment and increased affinity to these cells, and thus a greater proportion of this virus disseminates throughout the host via immune cells to the spleen and other organs. MNV-1.CW1 T301, on the other hand, has lower binding affinity and infectivity to these cells, and thus more virus stays proximal to the gastrointestinal tract. It is also possible that similar amounts of MNV-1.CW1 I301 and MNV-1.CW1 T301 disseminates to the spleen, but then the MNV-1.CW1 I301 variant has increased replication once in splenic cells, due to the richness of immune cells present.

The binding of MNV to CD300lf is thought to be a low affinity, high avidity interaction that requires a network of hydrophilic and hydrophobic interactions (25). Each virion is thought to interact with multiple CD300lf receptors, forming clusters that increase avidity (25, 44). Our results are consistent with this hypothesis and also suggest that this high avidity interaction is dependent on membrane fluidity, with low temperature and the depletion of cholesterol resulting in decreased virus binding. These results build upon the established idea that cholesterol is required for MNV endocytosis and further implicates the importance of lipid rafts (36, 45), which contain both cholesterol and CD300lf, and are important in signal transduction (38, 46–48). Sphingolipid biosynthesis is required to induce a conformational change in CD300lf to allow MNV infection (48), and thus it can be postulated that sphingolipids may also be required alongside cholesterol to regulate membrane fluidity required for lipid raft formation (47, 48). As isoleucine at VP1 301 enhances binding to cells when in suspension, one interpretation of our data is that increased hydrophobic interactions between the VP1 I301 variant and CD300lf confers increased affinity with receptor clusters that form at cholesterol-rich areas of the membrane. A similar interaction also occurs in other viruses, such as influenza, with cholesterol inducing nano-clusters of the glycosphingolipid receptor to increase virus infectivity (49). Furthermore, previous work has shown that GCDCA and metal ions can induce MNV P domain conformational changes that increase receptor affinity (6, 29). Future investigations should therefore investigate whether these factors can have additive effects in MNV attachment to suspension cells.

The results described in our study suggests that I301 plays a physiologically important role. It must be noted, however, that a previous study suggested the T301I substitution is a tissue culture adaptation, with MNV-3 collected from mice faeces 56 days post-infection reverting to T301 (15), which we did not observe in our experiments. The difference in these findings may be explained by experimental variations such as the location of infection, time post-infection, or both.

During infection of the host, viral quasi-species may provide the VP1 sequence diversity to generate viral sub-populations that allow access to different host cell types and widen dissemination. Furthermore, these sub-populations may change over time, depending on host immune pressures. Indeed, the T301I substitution has been previously identified as one of three mutations that occurred in an MNV escape mutant, following the addition of a monoclonal antibody that targeted VP1 (50). Viral quasi-species evolution is likely to be of relevance to HNV pathogenesis and chronic infection. Chronic HNV infection can persist for years in immunocompromised patients, leading to dehydration and nutrient deficiencies that can lead to mortality (19, 20, 51). Evolutionary studies have shown that HNV amino acid mutations accumulate throughout the chronic infection period (52, 53), with most being in the VP1 P domain (53). These evolutionary changes have also been linked to immune evasion, which leads to the changes in antigenic epitope of the virus (54). However, data that utilises virus-like particles (VLPs) and bioinformatics are contradictory as to whether this can change receptor-binding interactions (54, 55). Our study demonstrates a mechanism by which virus capsid evolution can significantly affect tissue tropism and susceptibility by enhancing interaction with the host cell. It can be postulated that this mechanism may also be utilised by HNV to avoid immune detection and influence chronic infection. This hypothesis should be investigated further as current reverse genetics systems (56, 57) are improved to allow the study of HNV infectivity.

## Methods

### Cells and mice

BHK-21 cells (obtained from ATCC) and RAW264.7 cells (gifted by Ian Clarke, University of Southampton) were maintained as previously described (6), with adherently grown BV-2 cells (gifted by Ian Goodfellow, University of Cambridge), maintained using the same method. Suspension grown BV-2 cells (referred to as BV-2S cells) were cultured in spinner flasks by maintaining a viable density of 0.5-1×10^6^ cells/mL with media changes every 2 days. WEHI-231 cells (obtained from ATCC) were maintained as previously described (58). Cells were incubated at 37°C and 5% CO_2_.

Balb/c mice were purchased from Jackson Laboratories (Bar Harbor, ME) and housed under specific-pathogen-free (SPF) and MNV-free conditions in accordance with federal and university guidelines. The protocol was approved by the University of Michigan Committee on Use and Care of Animals (UCUCA protocol number PRO00010484). Mice were allowed to acclimate in the facility for 6 days prior to infection. Mice were infected via oral gavage with 3×10^5^ PFU of virus in 200 μL/mouse. Tissues were harvested at 12 hours post infection and processed for plaque assay as described (59).

### Plasmid constructs

The plasmid, pT7-MNV*, containing the infectious clone sequence from MNV-1 strain CW1P3 (36) under control of T7 promoter was used for virus recovery. To exchange VP1 T301 standard two-step overlapping PCR mutagenesis was used with this plasmid as template (31). The pcDNA3.1(+)IRES GFP plasmid used for transfection experiments (kindly donated by Jamel Mankouri, University of Leeds) has already been described (61). Sequences of plasmids and primers are available on request.

### *In vitro* transcription and virus recovery

MNV plasmids were linearised with *Not*I and phenol/chloroform extracted before being used for *in vitro* transcription using the HiScribe™ T7 ARCA mRNA Kit (NEB), following the manufacturer’s instructions. RNA was purified and concentrated using the RNA Clean and Concentrator Kit (Zymo). RNA/DNA transfection was carried out as previously described (62). GFP fluorescence at 24 and 48 hours post transfection was analysed via the Incucyte S3 machine (Sartorius).

### TCID_50_ assay

Viral infectivity was determined using a TCID_50_ assay modified from Hwang et al (29), as per (6). For adherent TCID_50_ assays, BV-2 cells were seeded into 96 well plates at 2×10^4^ cells/well and left overnight before infection. For suspension TCID_50_ assays, viral dilutions were prepared and added to the plates first, prior to infection. TCID_50_ values were calculated according to the Spearman and Kärber algorithm (30).

### MTS assay

Cell viability in WEHI-231 cells was calculated 48 hours after MNV infection via the CellTiter 96® AQueous One Solution Cell Proliferation Assay kit, following manufacturer’s instructions. Absorbance was read on the Infinite F50 (Tecan) machine. Cytopathic effect was assigned to wells with values under 1. The number of positive wells was then used to calculate TCID_50_ values.

### Plaque assay

The plaque assay was performed from virus isolated from mouse tissue as previously described (59, 63). Data were normalized to the tissue weight and expressed as PFU per gram of tissue.

### MNV RNA extraction and sequencing

Viral RNA was extracted using the Direct-zol RNA miniprep kit (Zymo Research) according to the manufacturer’s instructions. For VP1 sequencing, ORF2 was amplified by RT-PCR using Superscript IV (Invitrogen), followed by second strand synthesis using Phusion DNA Polymerase (NEB). The sequence of the amplicon was determined by Sanger sequencing (Azenta). Sequences of primers used are available on request.

### One-step RT-qPCR

Virus sample was treated with 25 U/mL recombinant HS-Nuclease (MoBiTec) at 37□C for 30 minutes and viral RNA was extracted as previously described (62). RNA reverse transcription and cDNA amplification was then carried out by the GoTaq 1-Step RT-qPCR System (Promega), using established primers (31). CT values were converted to RNA copies/mL by analysing against a pT7-MNV* RNA standard curve of known values. The results were read using the Stratagene Mx3005P qPCR machine (Agilent Technologies).

### Western blot

SDS-PAGE and western blot analysis was carried out as previously described (27). Primary antibodies used were anti-MNV VP1 monoclonal antibody (MABF2097, Sigma-Aldrich), anti-CD300lf monoclonal (MAB27881, R&D Systems), anti-GAPDH monocolonal (60004-1, ProteinTech) and anti-BSA monoclonal antibody (66201-1, ProteinTech). Polyclonal anti-mouse (PA1-84388, Invitrogen) and anti-rabbit (HAF008, R&D Systems) HRP conjugates were employed as a secondary antibody. Blots were analysed on the G:BOX Chemi XX6 machine (Syngene).

### Flow cytometry

Detached adherent BV-2 cells or BV-2S cells (2×10^6^/mL) were analysed for CD300lf expression using a flow cytometry protocol previously described (64), with anti-CD300lf primary antibody (MAB27881, R&D Systems) and Alexafluor647 goat Anti-mouse IgG (A-21235, Invitrogen). The samples were analysed on a Cytoflex S flow cytometer (Beckman Coulter).

### Viral binding assay

Viral binding affinity to BV-S cells was determined using a binding assay modified from Berry and Tse (33). 10^5^ BV-2S cells were pre-treated with 50 μM dynasore (Ds; Cambridge Bioscience), 2 mM MβCD (Sigma-Aldrich) or 40 mM Los (Sigma-Aldrich) for 30 minutes at 37□C. MNV was added to the cells at an MOI of 1 and incubated at either 0□C or 37□C for 2 hours, before completing the binding assay as described.

### Statistics

Data were analysed and presented via GraphPad Prism v9.0 as mean ± SEM, N; biological repeat, with number of repeats stipulated in the figure legends. Statistical tests performed are also detailed within the figure legends with significant differences indicated by *p < 0.05; **p < 0.01; ***p < 0.001.

## Author Contributions

Conceptualization: Jake T. Mills, Susanna C. Minogue, Joseph S. Snowden, David J. Rowlands, Nicola J. Stonehouse, Christiane E. Wobus and Morgan R. Herod.

Investigation: Jake T. Mills, Susanna C. Minogue, Joseph S. Snowden, Wynter K.C. Arden, and Morgan R. Herod.

Supervision: David J. Rowlands, Christiane E. Wobus and Morgan R. Herod

Writing – original draft: Jake T. Mills and Morgan R. Herod.

Writing – review & editing: Jake T. Mills, David J. Rowlands, Nicola J. Stonehouse, Christiane E. Wobus and Morgan R. Herod.

## Acknowledgements

We thank Ian Goodfellow (University of Cambridge) and Ian Clarke (University of Southampton) for the murine cell lines.

