## Supplemental Figure 1 for "Amino acid substitutions in norovirus VP1 dictate cell tropism via an attachment process dependent on membrane mobility"

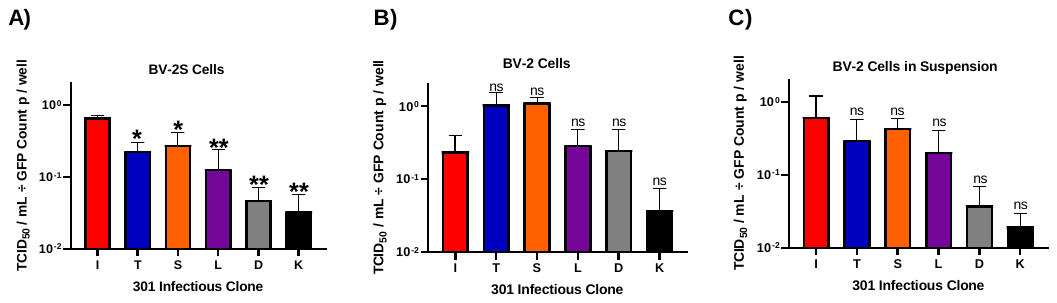
**Supplemental Figure 1: MNV I301 virus is preferentially recovered from BV-2S cells when normalised to GFP fluorescence.** MNV-1.CW1 infectious clone RNAs with the indicated amino acids at VP1 301 were transfected into BHK-21 cells alongside an IRES-GFP plasmid and virus-containing supernatants collected after 48 hours. Virus titre was determined by TCID_50_ assays on **(A)** BV-2S cells, **(B)** adherent BV-2 cells, or **(C)** adherent BV-2 cells infected in suspension. Data show mean TCID_50_/mL normalised to GFP fluorescence at 24 hours, with significance compared to I301 using one-way ANOVA with corrections for multiple comparisons (n = 3 ± SEM, *p<0.05; **p<0.01).
