## Supplemental Figure 2 for "Amino acid substitutions in norovirus VP1 dictate cell tropism via an attachment process dependent on membrane mobility"

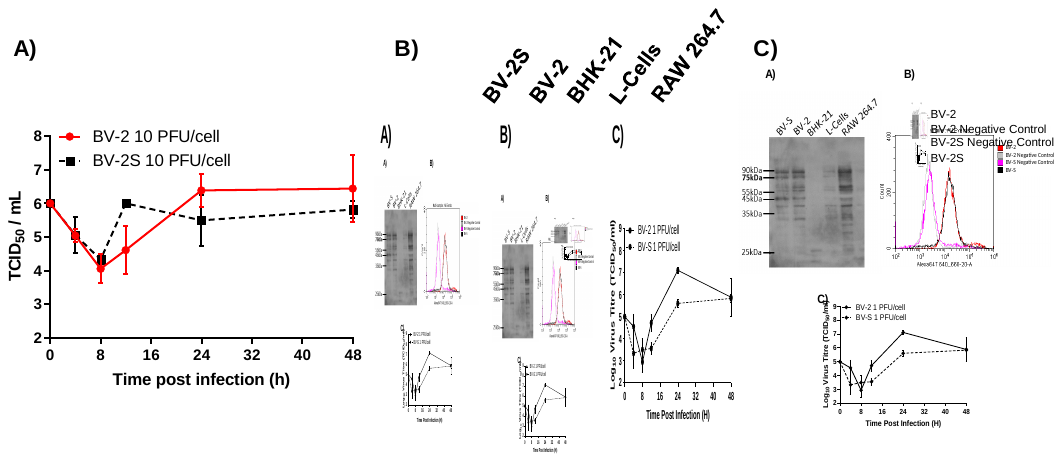
**Supplemental Figure 2: CD300lf expression is comparable in BV-2 cells and BV-2S cells. (A)** One-step growth curves of MNV-1.CW1 (T301) in BV-2 and BV-2S cells, infected at 10 PFU/cell. No significant differences were observed in titres between the cell types at each time point using the one-way ANOVA with corrections for multiple comparisons; data ± SEM, n = 3. CD300lf expression in the indicated cells was measured by **(B)** western blot (~75 kDa; n = 2; representative blot shown) and **(C)** flow cytometry (with negative no secondary antibody controls; n = 2; representative plot shown).
